# ISGylation-independent protection of cell growth by USP18 following interferon stimulation

**DOI:** 10.1101/2023.07.21.549904

**Authors:** Anne Clancy, Emma V. Rusilowicz-Jones, Iona Wallace, Kirby N. Swatek, Sylvie Urbé, Michael J. Clague

**Affiliations:** Biochemistry, Cell and Systems Biology, Institute of Systems, Molecular and Integrative Biology, University of Liverpool, Crown St., Liverpool, L69 3BX, UK; MRC Protein Phosphorylation and Ubiquitylation Unit, School of Life Sciences, University of Dundee, Dundee DD1 5EH, UK

**Keywords:** USP18, ISG15, ISGylation, interferon signaling, deubiquitylase

## Abstract

Type 1 interferon stimulation highly up-regulates all elements of a ubiquitin-like conjugation system that leads to ISGylation of target proteins. An ISG15-specific member of the deubiquitylase family, USP18, is up-regulated in a co-ordinated manner. USP18 can also provide a negative feedback by inhibiting JAK-STAT signaling through protein interactions independently of DUB activity. Here, we provide an acute example of this phenomenon, whereby the early expression of USP18, post-interferon treatment of HCT116 colon cancer cells is sufficient to fully suppress the expression of the ISG15 E1 enzyme, UBA7. Stimulation of lung adenocarcinoma A549 cells with interferon reduces their growth rate but they remain viable. In contrast, A549 USP18 knock-out cells show similar growth characteristics under basal conditions, but upon interferon stimulation a profound inhibition of cell growth is observed. We show that this contingency on USP18 is independent of ISGylation, suggesting non catalytic functions are required for viability. We also demonstrate that global deISGylation kinetics are very slow compared with deubiquitylation. This is not influenced by USP18 expression, suggesting that enhanced ISGylation in USP18 KO cells reflects increased conjugating activity.

## Introduction

Upon challenge with type 1 interferon, cells up-regulate hundreds of interferon-stimulated genes (ISGs) (1). Amongst these are a ubiquitin related molecule, ISG15 and enzymes which specifically mediate its conjugation; the E1 and E2 ligases UBA7 (UBE1L) and UBCH8 (UBE2L6) respectively (2-4). Ubiquitin-like modifications are counteracted by specific peptidases, which in the case of ubiquitin itself fall within a super-family of enzymes called deubiquitylases (DUBs) (5). The largest sub-family of DUBs are the ubiquitin specific proteases (USPs). Whilst these are largely promiscuous across ubiquitin chain linkage types, some have acquired a more stringent selectivity (5,6). For example CYLD shows specificity for linear and Lys63-linked ubiquitin chains (7,8). One member, USPL1, is dedicated to SUMO modifications rather than ubiquitin, whilst USP18 is highly selective for ISG15 (9-13). USP18 is also one of the most prominent ISGs (14).

Knock-out of USP18 leads to enhanced ISG15 conjugation. However, this cannot account for phenotypes observed in USP18^-/-^ mice, such as brain injury and early death. ISG15^-/-^ mice are healthy, but when both knock-outs are combined the USP18 associated defects dominate (15,16). USP18 negatively regulates JAK-STAT signaling independently of its isopeptidase activity by binding to the IFNAR2 sub-unit of the type 1 interferon receptor and interfering with the JAK receptor interaction (17,18). The JAK-STAT pathway is activated and prolonged in USP18^-/-^ cells following interferon stimulation leading to increased production of ISGs and greater resistance to cytopathic effects caused by a number of viral infections (19,20). Enhanced JAK-STAT signalling upon USP18 deletion in mice does not reflect a simple loss of isopeptidase activity and is independent of ISG15 conjugation, as it is retained by double knock-out mice for ISG15 or UBA7 (21).

The most prominent line of research regarding USP18 has been to investigate its role as part of the innate immune response to infection. Absence of USP18 leads to increased resistance to viral infections, but this effect is not always recapitulated by loss of enzymatic activity (22,23). There is also strong interest in USP18 as a potential therapeutic target in oncology (24). USP18 levels are elevated in diverse cancers and loss of USP18 has been show to be tumour suppressive in a mouse model of human mammary cancer (25,26). Interferon stimulation renders cell viability contingent on USP18 expression (19,27-29). There have been some contradictory reports as to whether catalytic activity is required for this effect, based on rescue experiments with catalytically inactive versions of USP18 (27,29). This is compounded by clear species differences in its biology (30). Here we have used an orthologous approach to resolve this controversy for human cell lines, thus clarifying appropriate strategies for therapeutic targeting.

The human genome contains seven E1 enzymes that are devoted to ubiquitin or related molecules. Inhibitors specific for UBE1 (ubiquitin, TAK-243) and UBA3 (NEDD8, MLN4924) have been characterised and entered clinical trials (31,32). No specific inhibitors have been described that target UBA7, the E1 specific to the ISGylation cascade (33). Here we have used acute application of a pan-E1 inhibitor, compound 1, to block all ubiquitin-like conjugation activities (34). This has allowed us to compare the kinetics of dissipation of the ISGylome with the ubiquitylome and to assess the role of USP18 in this regard.

## Results

### High basal expression and USP18 sensitivity in H1650 cells reflects constitutive STAT1 activation

By consulting global CRISPR/Cas9 screening data for cell viability across hundreds of cell lines, we observe that USP18 and ISG15 are highly correlated and anti-correlated with components of the interferon signaling system (35,36). A small number of cell lines are highly sensitive to both ISG15 or USP18 loss (Figure 1A). We speculated that this reflects the known loss of cell viability reported for a variety of cell lines when interferon stimulation and USP18 loss are combined. We confirmed that H1650 cells express high levels of USP18 under basal conditions, compared with other cell lines (Figure 1B). This is accompanied by elevated levels of ISG15 and ISGylation that mirror interferon stimulated lung adenocarcinoma cells (A549) and is likely due to constitutive STAT1 activation (Figure 1B,C). In these H1650 cells levels of ISGylation cannot be further enhanced by acute interferon stimulation. We next showed that siRNA depletion of USP18 in H1650 cells reduces cell viability in accordance with screening data (Figure 1D). We investigated why H1650 cells show constitutive activation of ISGylation. They carry an activating EGFR mutation, but 72 hour treatment with a selective EGFR inhibitor (Afatinib) had no effect on STAT1 phosphorylation status or expression of USP18 and ISG15 (Figure 1E). In fact constitutive ISGylation must represent autocrine activation in these cells as the application of H1650 conditioned media to A549 cells induces ISGylation in those cells in distinction to conditioned media derived from A549 or H1975 cells. (Figure 1F).

**Figure 1:**
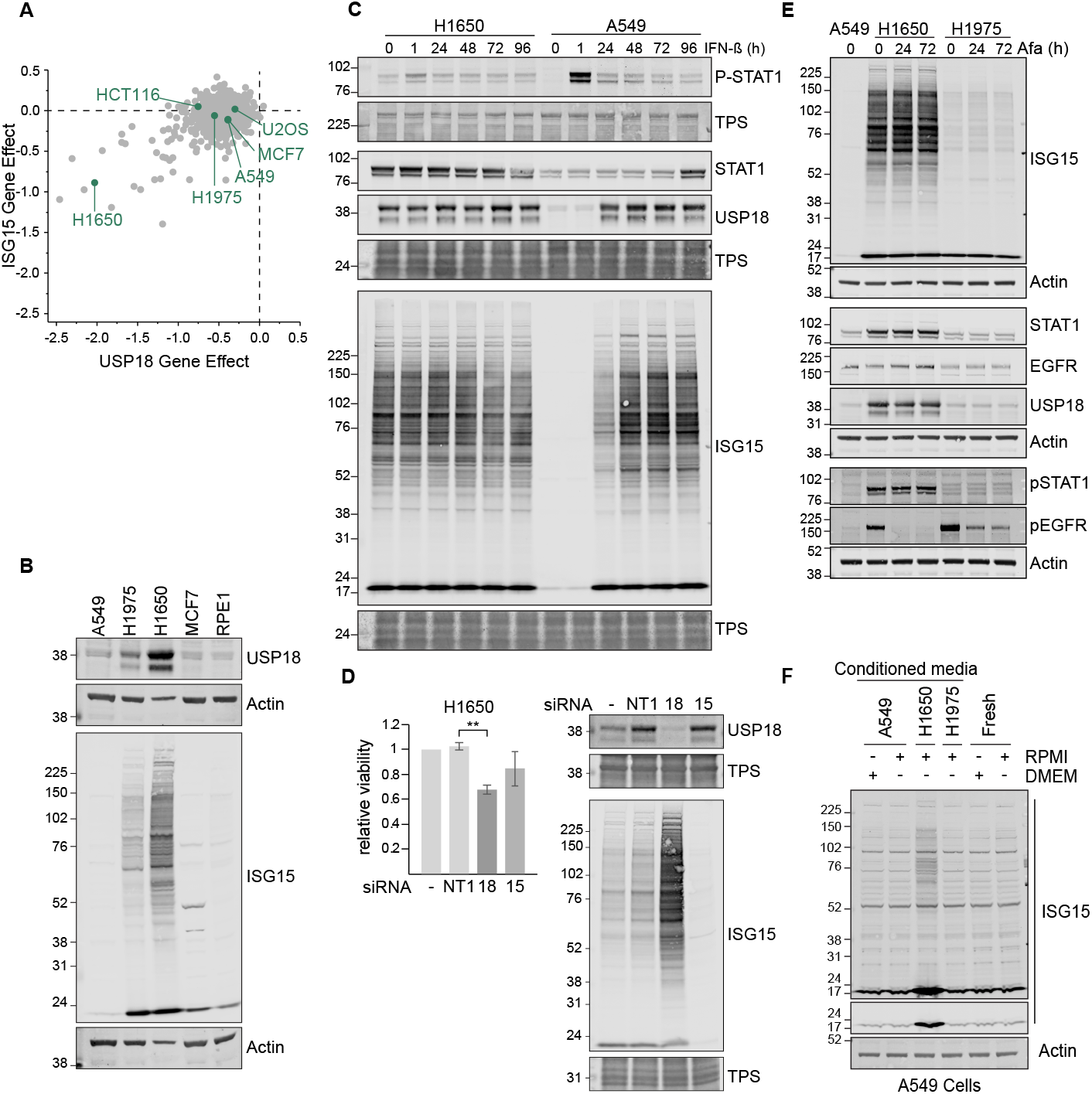
High constitutive levels of USP18 in H1650 cells reflect pSTAT1 status and render them sensitive to USP18 depletion. (A) Correlation of Gene Dependency Score (CRISPR (Depmap Public 23Q2+Score, Chronos)) across cell lines predicts H1650 cell line co-dependency on USP18 and ISG15. Data have been downloaded from the Depmap portal (36). (B) Comparison of USP18 and ISGylation levels across cell lines. (C) Contrasting responses of H1650 and A549 cells to IFN-ß stimulation (50pM), constitutive versus induced USP18 expression and ISGylation. TPS = total protein stain. (D) MTS assay for H1650 cell viability following siRNA mediated depletion of USP18 or ISG15 for 96 hours, n=3, error bars represent standard deviation, **p<0.01; one-way ANOVA with Tukey Multiple comparisons test. (E) H1650 cells subjected to EGFR inhibition with Afatinib (50 nM). (F) Media were conditioned on the indicated cells for 72 hours and then applied to A549 cells as indicated for 48 hours. Fresh: Unconditioned media.

### Differential response of the ISGylation cascade components to interferon reveals repression of UBA7 by USP18

We next surveyed six cell lines for their responsiveness to interferon with respect to ISGylation components USP18, UBA7 (E1) and the cognate E2, UBCH8. HCT116 colon cancer cells were distinct from other cell lines. USP18 and ISG15 expression is efficiently induced in these cells but UBA7 is not discernible and UBCH8 highly reduced (Figure 2A). A less stringent suppression of UBA7 is also seen with MCF7 breast cancer cells, which also express high levels of USP18 (Figure 2A). This renders HCT116 cells defective in up-regulating ISGylation despite intact interferon signaling. Consequently, depletion of USP18 restores interferon-dependent UBA7/ UBCH8 expression in HCT116 cells and ISGylation (Figure 2B).

**Figure 2:**
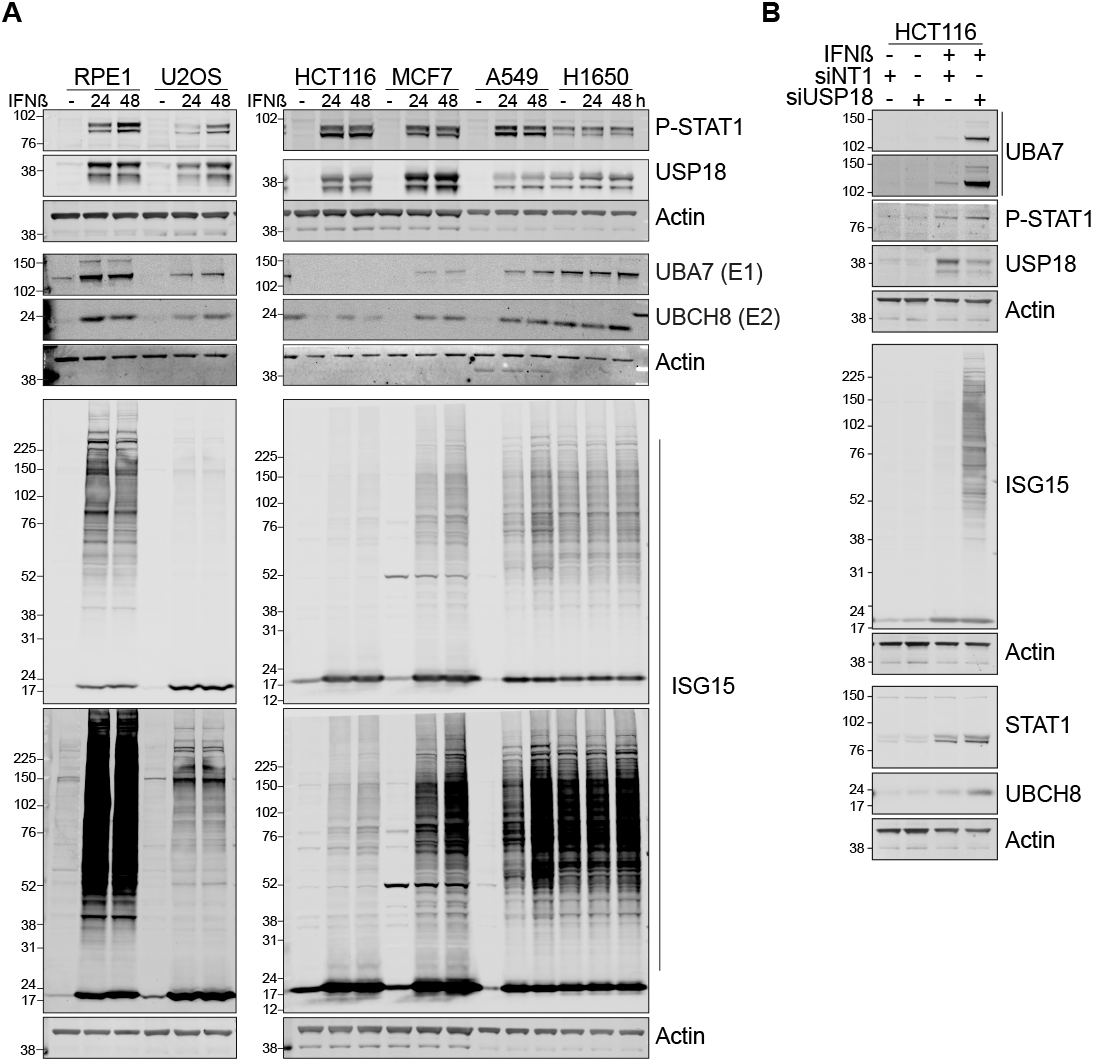
Differential responses to IFN-ß stimulation reveal USP18 dependent suppression of UBA7 expression and ISGylation in HCT116 cells. (A) IFN-ß induction of ISG15 conjugating enzymes, USP18 and ISGylation across multiple cell types (B) Depletion of USP18 by siRNA restores IFN-ß (50pM, 68 hours) induction of UBA7 and ISGylation in HCT116 cells.

### Interferon contingency is independent of UBA7 in USP18 KO cells

We used CRISPR-Cas9 gene editing to generate A549 USP18 knock-out (KO) cell clones, which show a characteristic up-regulation of pSTAT1 and ISGylation upon acute interferon stimulation (Figure 3A). Using an Incucyte imaging system to measure cell growth, we observe that interferon stimulation retards growth of wild type parental cells, but completely blocks any growth of USP18 KO cells (Figure 3B). A similarly dramatic effect is seen with an orthologous colony formation assay (Figure 3C and D). To ask if this might reflect the enhanced ISGylation, we depleted UBA7 in these cell lines using siRNA. Importantly, following interferon stimulation, the small amount of residual ISGylation in these UBA7-depleted cells is significantly below the levels of ISGylation in the parental cell lines, which exhibit cell growth under interferon stimulation (Figure 3E). Thus, if the effect of USP18 loss on cell viability is dependent on its suppression of ISGylation then cell growth should be restored by depletion of UBA7. Our data, which show no impact of UBA7 depletion on cell viability clearly discount this and instead support a model whereby USP18 protects from interferon independently of deISGylation activity (Figure 3F).

**Figure 3:**
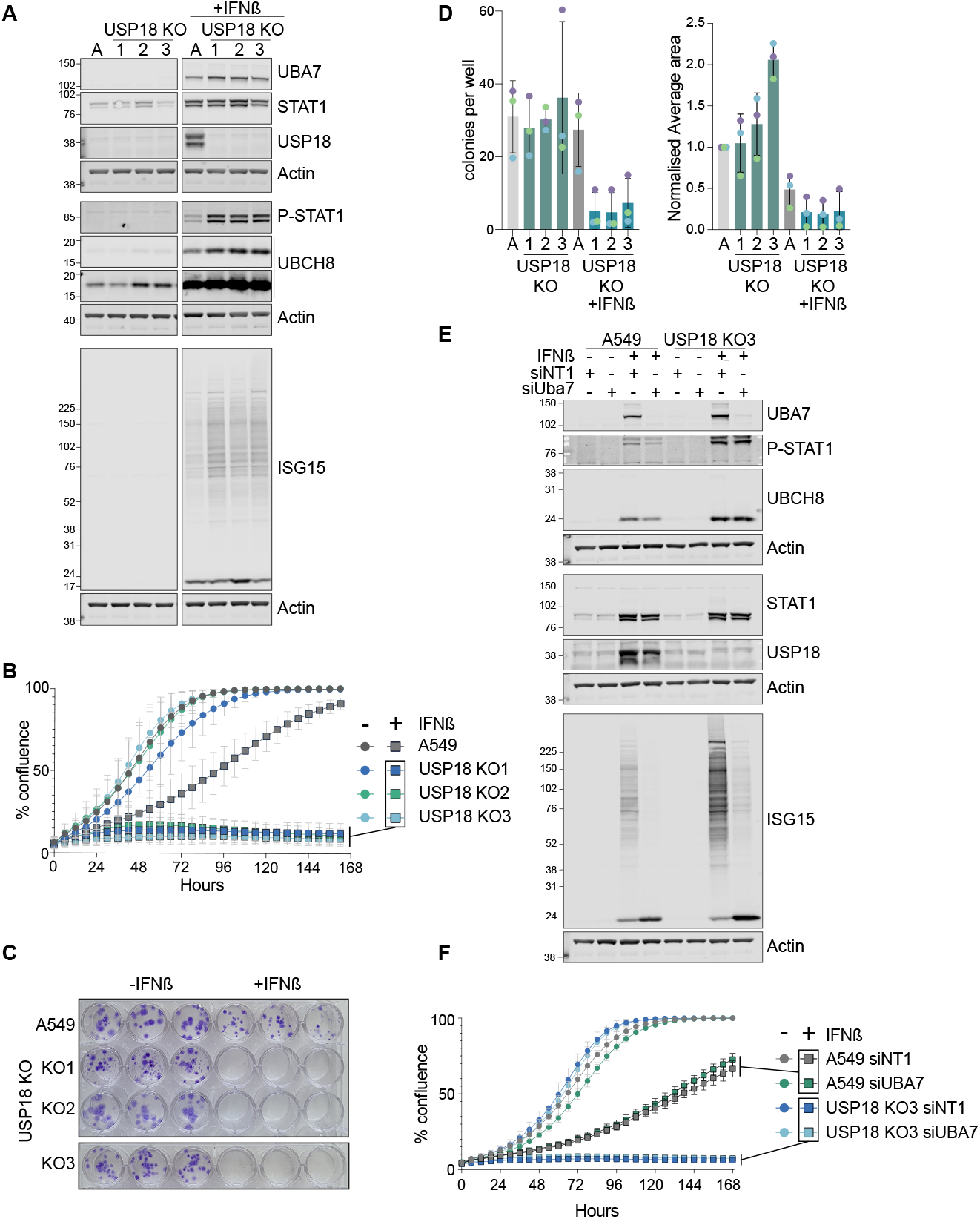
USP18 KO A549 cell growth is sensitive to IFN-ß stimulation independent of UBA7 expression and ISGylation capacity. (A) Characterisation of the IFNß response (50 pM, 24 hours) of 3 individual USP18 KO clones (1,2,3) response to IFN-ß, 50 pM for 24 hours. A= parental A549 cells. (B) Incucyte cell growth assay for parental and USP18 KO A549 cells ± IFN-ß (50 pM, 24 hours). (C) Colony formation assay for parental and USP18 KO A549 cells ± IFN-ß (50 pM) visualised after 14 days. (D) Quantitation of colony formation assays, n=3, error bars represent standard deviation. (E) Effect of siRNA-mediated depletion (72 hours) of UBA7 on ISGylation in parental and USP18 KO A549 cells, (IFNß 50 pM, 68 hours). (F) Incucyte cell growth assay for parental and USP18 KO A549 cells ± IFN-ß and ±UBA7.

### ISGylation is a highly stable PTM

With isogenic A549 cells ±USP18 in hand, we designed an experiment to determine its effects on the global turnover of ISGylation, first by accumulating ISGylation with interferon stimulation and subsequently applying a pan-E1 inhibitor, compound 1, whilst withdrawing interferon (Figure 4A,B) (37). To limit complications owing to toxicity, we have then monitored the decay of global ISGylation and ubiquitylation for a relatively short time period of 6 hours (Figure 4A). We adjusted the loading of samples on the gels to reflect roughly equal levels of ISGylation at time zero (4 times more sample for parental compared to USP18 KO) or ubiquitylation (equal protein loaded). The adjusted loading also makes it easier to visualise differences in banding patterns on ISG15 blots between the parental and knock-out cells. In setting up this experiment we hoped to be able to estimate the contribution of USP18 to the deISGylation kinetics. However to our surprise the level of ISGylation remained constant across all identifiable bands over this 6 hour chase interval (Figure 4B). Deubiquitylation is striking and rapid, complete within 2 hours, similar to previous reports using a specific UBA1 inhibitor (38), confirming compound activity on intact cells. The compound is clearly active on the ubiquitin and ISG15 conjugation cascade as evidenced by pronounced deubiquitylation and discharge of the thioester linked ISG15 from UBA7 and ubiquitin from the E2 enzyme UBC13, revealed on non-reducing gels by the increase in the lower molecular weight uncharged forms (Figure 4C). This was further confirmed with an *in vitro* assay using purified UBA7 and UBCH8, where application of the pan-E1 inhibitor leads to loss of ISG15-charged enzymes (Figure 4D). Thus in A549 cells, ISGylation is a highly stable post-translational modification even when USP18 expression has been induced by 24 hour interferon stimulation. This indicates that the increased levels of ISGylation in USP18 KO cells reflects increased conjugation rather than decreased deISGylation activity, in accord with increased UBA7 and UBCH8 levels (Figure 2). Any conceivable alternative interpretation of our results would require that there is a residual ISG15-conjugating activity that exactly balances deconjugating activity, which we think is unlikely.

**Figure 4:**
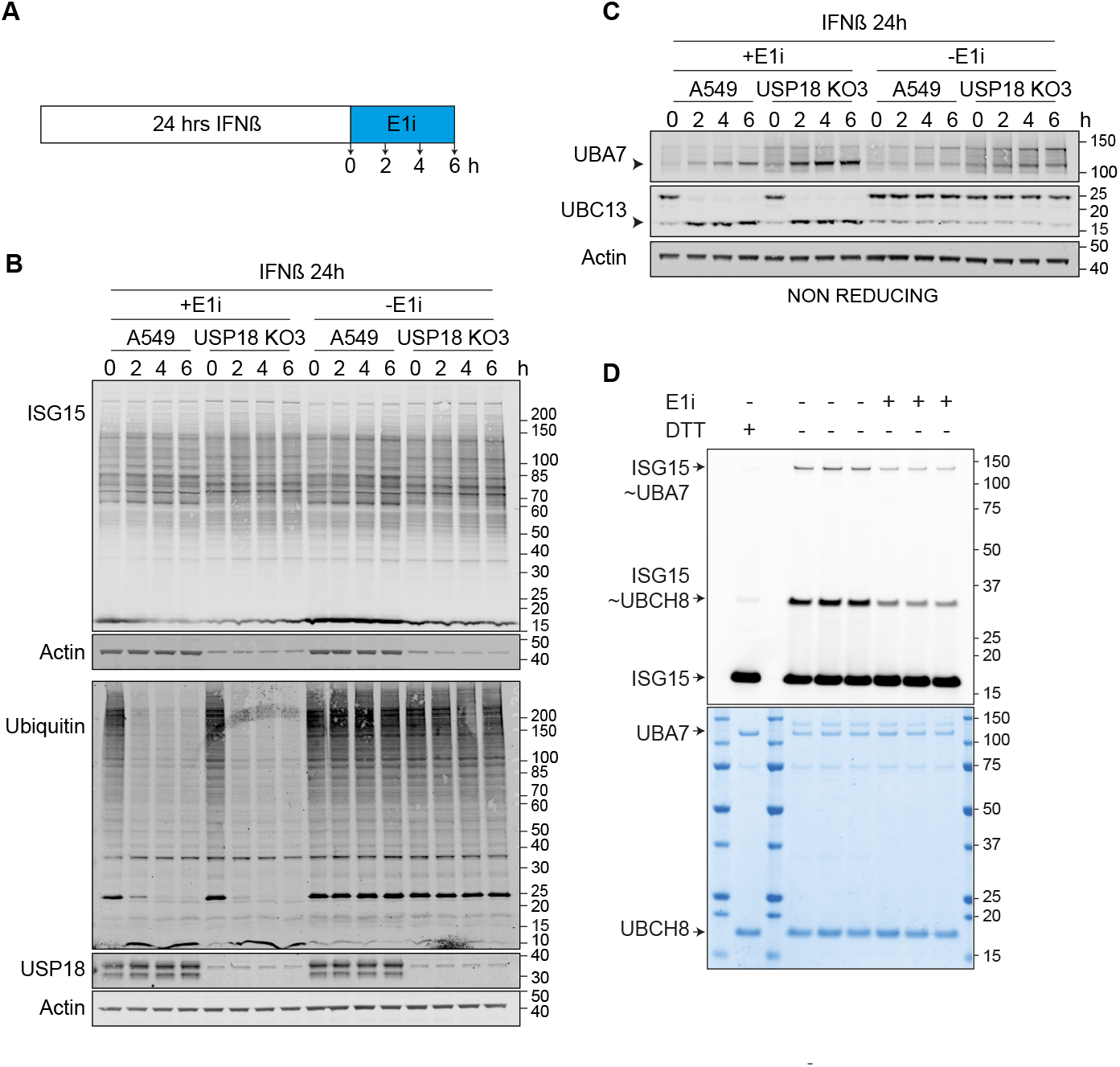
Global deISGylation kinetics. (A-C) A549 WT and USP18 KO cells were treated with 50 pM IFNß for 24 hours to accumulate ISGylated proteins. IFNß was then removed and cells were treated with a pan E1 inhibitor, compound 1 (E1i, 25 μM) for the indicated times. Samples were analysed on (B) reducing and (C) non-reducing gels. For analysing ISGylation, 4 times less protein was loaded for USP18 KO samples compared to A549 WT in order to equalise starting levels of ISG15. (D) E1 enzyme inhibition assay. The E1 inhibitor (E1i) was first added to preformed UBA7∼ISG15 conjugates (using fluorescently labeled ISG15) and then UBCH8 enzyme was added to the reaction. E1 and E2 charging was monitored by fluorescent imaging of SDS-PAGE gels (upper panel, coomassie stain lower panel), ∼ denotes a thioester bond.

## Discussion

The contingency of cell growth/survival on USP18 expression under interferon stimulatory conditions offers a therapeutic opportunity. Many tumour types show elevated expression of interferon responsive genes including USP18 relative to neighbouring tissue (24). With the advancement of specific small molecule inhibitors of USP family members (39-43), USP18 has attracted considerable attention as a potential small molecule target. To clarify any future drug development strategy, we set out to resolve conflicting reports as to whether USP18 catalytic activity is required for its pro-survival effect. Rather than use a rescue approach with wild-type versus catalytically inactive proteins, we have adopted an alternative strategy of suppressing ISGylation in A549 cells and showing that a dependency upon USP18 expression is 100% maintained. This concurs with previous findings in mouse models (21). It implies that the principal effect of USP18 relevant to cell growth is attenuation of JAK/STAT signaling in a catalytically independent manner. Thus our findings support the case for USP18 as a therapeutic target, but suggest that the appropriate strategy will be to develop PROTAC molecules rather than catalytic inhibitors. In fact such inhibitors could be developed as PROTACs.

HCT116 colon cancer cells provide a striking illustration of the profound negative feedback effect exerted by USP18. In these cells, USP18 is expressed following interferon stimulation, yet in distinction to other cell types, there is minimal induction of ISGylation, UBA7 and UBCH8. We show that it is the presence of USP18, rather than an intrinsic unresponsiveness to interferon of these cells, that prevents this expression, which is restored following USP18 depletion. We propose that differences in timing of expression can affect the outcome between different cell types. If USP18 is able to accumulate ahead of other ISG-responsive proteins, then it can effectively suppress the entire ISGylation cascade.

With isogenic USP18 KO cells in hand, we assumed that we would have one of the necessary tools to assess the contribution of USP18 to the cellular dynamics of deISGylation. The other tool required is a means to acutely switch off ISGylation following the accumulation of a global ISGylation signal and significant levels of USP18 following interferon stimulation. For ubiquitylation this is achieved by the application of a specific UBA1 inhibitor i.e. TAK-243. No such specific inhibitor for UBA7 exists to our knowledge and we have therefore turned to using a pan-E1 inhibitor. This led us to our third major result; the surprising observation that once attached, ISGylation is a very stable post-translational modification, with no evidence of decay over a six hour period. This observation rendered this line of enquiry into USP18 moot, but further serves to emphasise the pre-eminence of non-catalytic roles of USP18.

## Materials and Methods

### Cell culture

A549, U2OS and MCF7 cells were cultured in DMEM with GlutaMAX+ 10% FBS, H1650 and H1975 cells in RPMI + 10% FBS, HCT116 in McCoys media + 10% FBS and RPE1 cells in DMEM/F12 media + 10% FBS, at 37^°^C and 5% CO^2^. Cells were routinely checked for mycoplasma. IFNß (Peprotech) was made up in PBS/1%BSA and added to media at 50pM. Compound 1 (Merck/Astatech) was made up in DMSO and added to media at 25*µ*M. For siRNA experiments, cells were treated with 20 nM of non-targeting (NT1) or target-specific siRNA oligonucleotides (Dharmacon On-Target Plus), using Lipofectamine RNAi-MAX (Invitrogen) according to manufacturer’s instructions. Medium was exchanged 24 h after transfection.

### MTS assay

CellTiter 96® AQueous One Solution Cell Proliferation Assay, Promega. Cells were grown in 96 well plates, 100 *µ*l per well, 20 *µ*l of assay reagent was added and incubated for 2-4h at 37^°^C and 5% CO_2_, subsequently absorbance was read at 490 nM.

### Colony Assay

Cells were seeded in a 24 well plate at 20-25 cells per well, 24 hours later relevant wells were treated with 50 pM IFNß. After 8-14 days of growth, cells were fixed in methanol at -20°C and stained with 1mg/ml crystal violet, in 20% methanol. Plates were photographed and scanned (GelCount, Oxford Optronix), area and colonies per well were calculated with GelCount software.

### Incucyte Growth Assay

Cells were grown in the presence or absence of 50 pM IFNß for 24 h, then replated at 3000 cells/cm^2^ in 96 well plates in triplicate. 4 frames per well were imaged every 6 h, 10x objective in an Incucyte (Essen Biosciences) at 37^°^C and 5% CO_2_. Percentage confluence was calculated with Incucyte analysis software.

### USP18 KO cell generation

Knock-out cells were generated using CRISPR-Cas9 technology with three independent USP18 specific sgRNAs targeting exon 3 (sgRNA1: TCAGGTGTTCGTAATGAATG, sgRNA2: AATCAAGGAGTTAAGGCAGC, sgRNA3: CTGGTTGGTTTACACAACAT). These sgRNAs were cloned into the pSpCas9(BB)-2A-GFP (PX458) vector (a gift from Feng Zhang, Addgene plasmid #48138) and transfected into A549 cells. GFP-positive cells were FACS sorted 24 h after transfection and single cell diluted. Individual clones were amplified and characterised by Western blotting and genomic DNA sequencing. Clone 3 was selected for use in further experiments.

### Lysis and Western Blot analysis

Cultured cells were lysed for 10 minutes on ice in RIPA buffer (10 mM Tris–HCl pH 7.5, 150 mM NaCl, 1% Triton X-100, 0.1% SDS, 1% sodium deoxycholate) supplemented with mammalian protease inhibitor cocktail and PhosSTOP (SIGMA). Proteins were resolved using SDS–PAGE (Invitrogen NuPage gel 4–12%), transferred to nitrocellulose membrane, blocked in 5% fat-free milk or 5% bovine serum albumin in TBS supplemented with Tween-20, and probed with primary antibodies. Visualisation and quantification of Western blots were performed using IRdye 800CW and 680LT coupled secondary antibodies and an Odyssey infrared scanner (LI-COR Biosciences, Lincoln, NE).

### In vitro ISG15 charging assay

E1 charging reactions were performed by mixing 3 *µ*M fluorescent ISG15 with 2.5 *µ*M E1 enzyme (UBA7) for 15 minutes in reaction buffer (50 mM HEPES pH 7.5, 150 mM NaCl, 5 mM MgCl2, 5 mM ATP). Reactions were stopped by diluting the sample 10-fold in quenching buffer (50 mM HEPES pH 7.5, 150 mM NaCl, 100 mM EDTA). DMSO (1%) or the E1 inhibitor (25 *µ*M) was then added to the quenched E1 charging reactions and incubated for 20 minutes. To assess E1 inhibition, 2 *µ*M E2 enzyme (UBCH8) was added for 15 minutes and reactions were stopped in 4X LDS sample buffer (-/+ DTT, as indicated). The samples were resolved by SDS-PAGE gels and E1/E2 charging was visualised by fluorescent imaging.

